# Meiotic Sex Chromosome Inactivation: conservation across the *Drosophila* genus

**DOI:** 10.1101/2024.12.04.626848

**Authors:** Camila C. Avelino, Carolina A. Mendonca, Gabriel Goldstein, Carvalho A. Bernardo, Maria D. Vibranovski

**Affiliations:** Department of Genetics and Evolutionary Biology, Institute of Biosciences, University of São Paulo, São Paulo, SP, Brazil; School of Pharmaceutical Sciences, University of São Paulo, São Paulo, SP, Brazil; Departamento de Genética, Universidade Federal do Rio de Janeiro, RJ, Brazil

## Abstract

The inherent difference between sex chromosomes in males and females drives conflicts in gene expression, leading to adaptations such as Meiotic Sex Chromosome Inactivation (MSCI). In this study, we explore the evolutionary dynamics of MSCI within the *Drosophila* genus by analyzing transcriptomes across different stages of spermatogenesis in *D. melanogaster* and its progressively more distant relatives, *D. simulans*, *D. willistoni*, and *D. mojavensis*. Stage-specific bulk RNA sequencing, showing a strong correlation in spermatogenic gene expression patterns among these species, revealed that MSCI dates back to the early evolution of the *Drosophila* genus, impacting the regulation of both coding and long non-coding RNAs. Notably, for newly evolved genes, X-linked genes show higher expression levels than autosomal genes during mitosis and meiosis, indicating that MSCI predominantly regulates older genes. In contrast, newly evolved autosomal genes exhibit a gradual increase in expression throughout spermatogenesis, reaching their peak in the post-meiotic phase. During this phase, the expression of X-linked new genes decreases, eventually aligning with that of autosomal genes. This expression pattern suggests that haploid selection plays a crucial role in the regulation of new genes, with monoallelic expression of the X chromosome providing an advantage across all stages of germline development, while autosomal gene expression gains a selective edge primarily in the post-meiotic phase. Together, these findings provide new insights into the evolution of sex chromosomes and highlight the critical role of MSCI in shaping gene expression profiles in *Drosophila*.

## Introduction

The occurrence of sex chromosome systems across phylogenetically diverse groups implies that various taxa have been subjected to analogous selective forces in the evolution of sex chromosomes that have arisen independently (Charlesworth, 1996). A remarkable feature of sex chromosome evolution is the phenomenon of Meiotic Sex Chromosome Inactivation (MSCI) (Lifschytz & Lindsley, 1972; Solari, 1974; Turner, 2007). During male meiosis, the X chromosome undergoes transcriptional silencing, a process that has been documented across diverse taxonomic groups, including mammals (Richler et al., 1992), grasshoppers (Cabrero et al., 2007), nematodes (Kelly et al., 2002; Daish & Grützer, 2010), and *Drosophila* (Hense et al., 2007; Vibranovski et al., 2009; Mahadevaraju et al., 2021). As these sex-chromosome systems evolved independently (Charlesworth, 1996), the repeated occurrence of MSCI suggests that it has a major role in sex chromosome dynamics (Daish & Grützer, 2010; Daish & Grützer, 2019).

In *Drosophila*, MSCI is driven by the absence of active RNA polymerase II (RNA pol II-S2) on the X chromosome territory in spermatocytes (Mahadevaraju et al., 2021). MSCI has primarily been studied in *D. melanogaster*, leaving open questions about when this silencing mechanism originated and its consequences for genes moving to/from the sex chromosomes after MSCI evolution.

To gain a comprehensive understanding of MSCI evolution within the *Drosophila* genus, we conducted stage-specific RNA-seq across the main stages of spermatogenesis (mitosis, meiosis, and post-meiosis) in four species: *D. melanogaster*, *D. simulans*, *D. willistoni*, and *D. mojavensis*. Our adapted dissection method reliably captured comparable expression profiles from spermatogenesis stages across species, revealing a conserved MSCI-associated downregulation on the X chromosome, including *D. willistoni*’s neo-X chromosome. Notably, MSCI extends to long noncoding RNAs (lncRNAs) in *D. melanogaster*, and an expression signature remains on the ancestral X chromosome (dot chromosome) across all species, although this signal appears to be diminished over time.

Our RNA-seq method, also optimized for profiling post-meiotic cells in four *Drosophila* species, provided an in-depth view of spermatogenic expression dynamics, allowing for a comprehensive analysis of gene expression patterns across stages and their relationships with gene age. This approach enabled us to examine the effects of MSCI and the role of haploid selection on new genes. Notably, MSCI does not impact newly emerged X-linked genes in *D. melanogaster*, which retain higher expression levels throughout the post-meiotic stages. This suggests that MSCI may require time to affect new sequences and underscores the advantage of exposing adaptive alleles in novel phenotypes influenced by single-copy chromosome expression.

## Results

### Data and method reproducibility across species

Given the temporal distribution of *Drosophila* spermatogenesis along the testis (Fuller, 1993), dissections based on Vibranovski et al. (2009) methods (Figure 1-A) were undertaken to isolate cells from the main stages of spermatogenesis. This technique relies on cellular morphology as a guide to isolate regions enriched for three major phases of *D. melanogaster* spermatogenesis: mitosis, meiosis, and post-meiosis (Figure 1-A).

**Figure 1.**
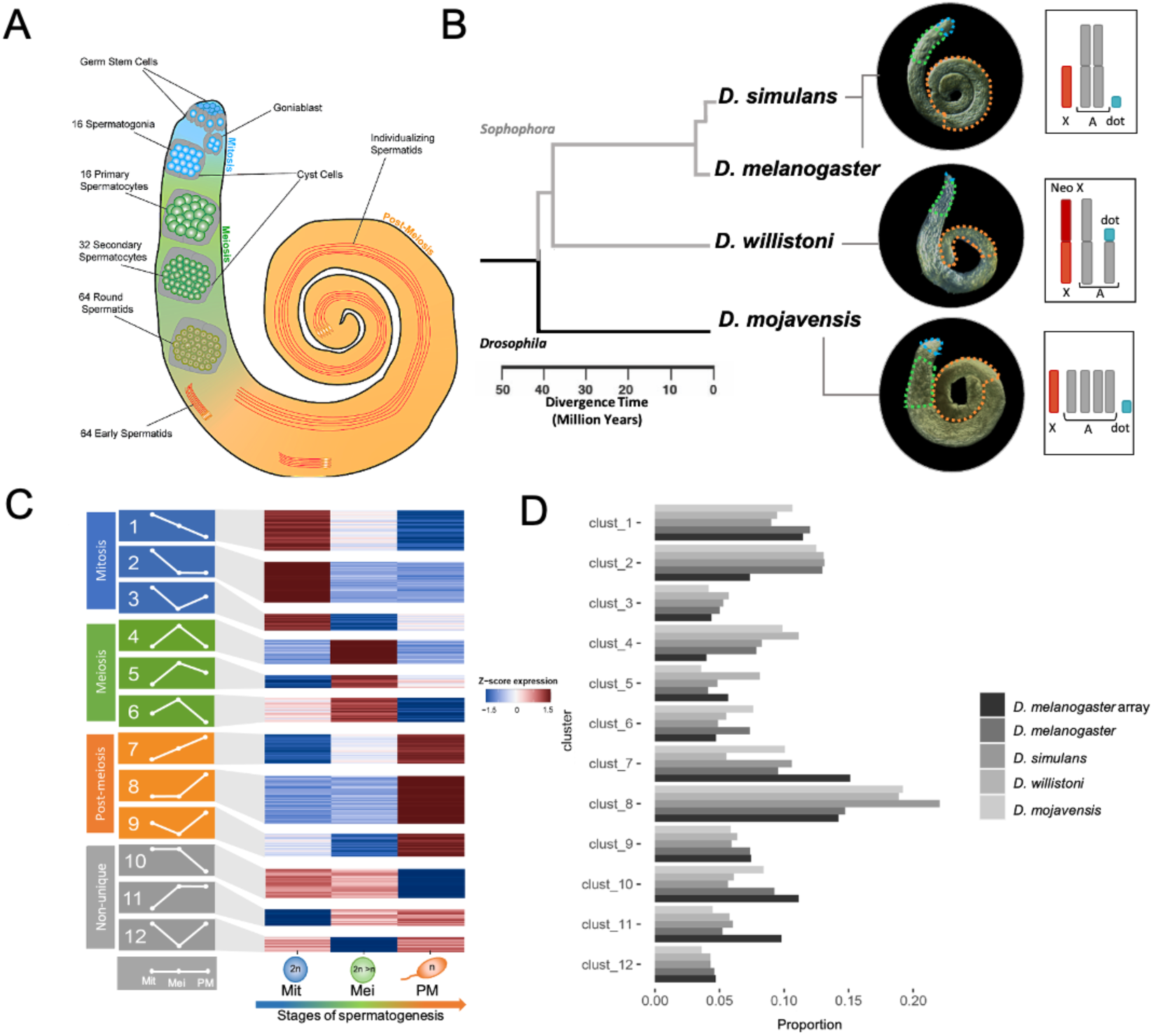
Stage-Specific Transcriptional Profiling During *Drosophila* Spermatogenesis. **A)** Main cell types involved in *Drosophila* spermatogenesis and their spatial localization within the testis. **B)** Phylogenetic tree of selected *Drosophila* species analyzed in this study, accompanied by whole testis images highlighting regions enriched with mitotic (blue), meiotic (green), and post-meiotic (orange) cells. Chromosome insets depict the karyotypes for each species, marking autosomes (gray), X chromosome (red), Neo-X chromosome (dark red), and dot chromosome (cyan). Notably, *D. willistoni* exhibits two chromosomal fusions involving the X and dot chromosomes with autosomes. **C)** Heatmap displaying k-means clustering of gene expression (Z-scores) across different stages of spermatogenesis: mitosis (Mit: blue), meiosis (Mei: green), and post-meiosis (PM: orange). Gray indicates non-unique profiles with broad expressions. **D)** Comparison of transcriptome cluster proportions between microarray data (Vibranovski et al., 2009) and RNA-seq data from this study across four species.

Cell enrichment isolation for RNA sequencing was conducted across four species within the *Drosophila* genus: *D. melanogaster*, *D. simulans*, recognized as a cryptic species; *D. willistoni*, examined as a sister group, featuring a neo-X chromosome (Tamura et al., 2004; Richards et al., 2005); and lastly, *D. mojavensis*, serving as an outgroup to the *Sophophora* subgenus (Figure 1-B). Since the technique was initially standardized in *D. melanogaster* (Vibranovski et al., 2009), we adapted it to the other species under scrutiny to ensure the appropriate section of each enriched region (See Materials and Methods; Figure S1).

First, our stage-specific *D. melanogaster* transcriptome from RNA-seq strongly correlates with previous data obtained using microarrays (Pearson correlation > 0.85; Figure S2). Additionally, high reproducibility was achieved across biological replicates for all spermatogenesis stages (> 90%; Figure S3). Moreover, samples from the same stage cluster closely in a within-species Principal Component Analysis (PCA), clearly separating from samples at other stages (Figure S4-A).

To facilitate interspecies comparisons, we needed to ensure that the adaptations applied to the dissection method would accurately capture equivalent spermatogenesis stages across species. For further analyses, orthologs following stringent criteria (1,913 out of 6,110 genes) were selected: (1) genes with only one annotated transcript (enabling 1:1 comparisons), (2) genes on the same Müller element, and (3) excluding those located on the Y chromosome and the D element (which functions as a Neo-X chromosome in *D. willistoni* and an autosome in other species).

Firstly, the PCA analysis in Figure S4-B examines inter- and intra-species variation in *Drosophila* gene expression during spermatogenesis. It reveals that the variance between spermatogenesis stages (PC1, 35.2%) is greater than the variance between species within the same stage (PC2, 14.9%), indicating that gene expression profiles are more similar across species at the same stage than across different stages. Additionally, the selected gene set exhibited a consistent high correlation across species and stages (Pearson correlation > 0.62, Figure S5, Table S5). Finally, in alignment with a prior investigation for *Drosophila* microarray data (Vibranovski et al. 2009, Raíces et al., 2019), our findings unveiled a total of 12 distinct expression profile clusters across all transcriptomes (Figure 1-C). Notably, the proportions of these clusters within each species demonstrated substantial similarity across transcriptomes (Figure 1D). Altogether, these results highlight the reproducibility and inter-species applicability of the dissection method originally established in *D. melanogaster* by Vibranovski et al. (2009), despite potential variations in cell proportions between species and expression detection methods.

### MSCI across the Drosophila genus

To examine the MSCI-induced downregulation of X-linked genes relative to autosomes, we performed a comparative analysis of the proportion of gene products differentially expressed between meiosis and mitosis. Consistent with prior microarray findings in *D. melanogaster*, our RNA-seq data revealed a higher proportion of underexpressed coding sequences on the X chromosome during meiosis compared to autosomes across all species—a key indicator of MSCI in *Drosophila* (Figure 2-B). Conversely, as an indirect effect of MSCI, autosomes showed a significantly higher proportion of overexpressed genes during meiosis compared to the X chromosome across all species (Figure 2-B).

**Figure 2.**
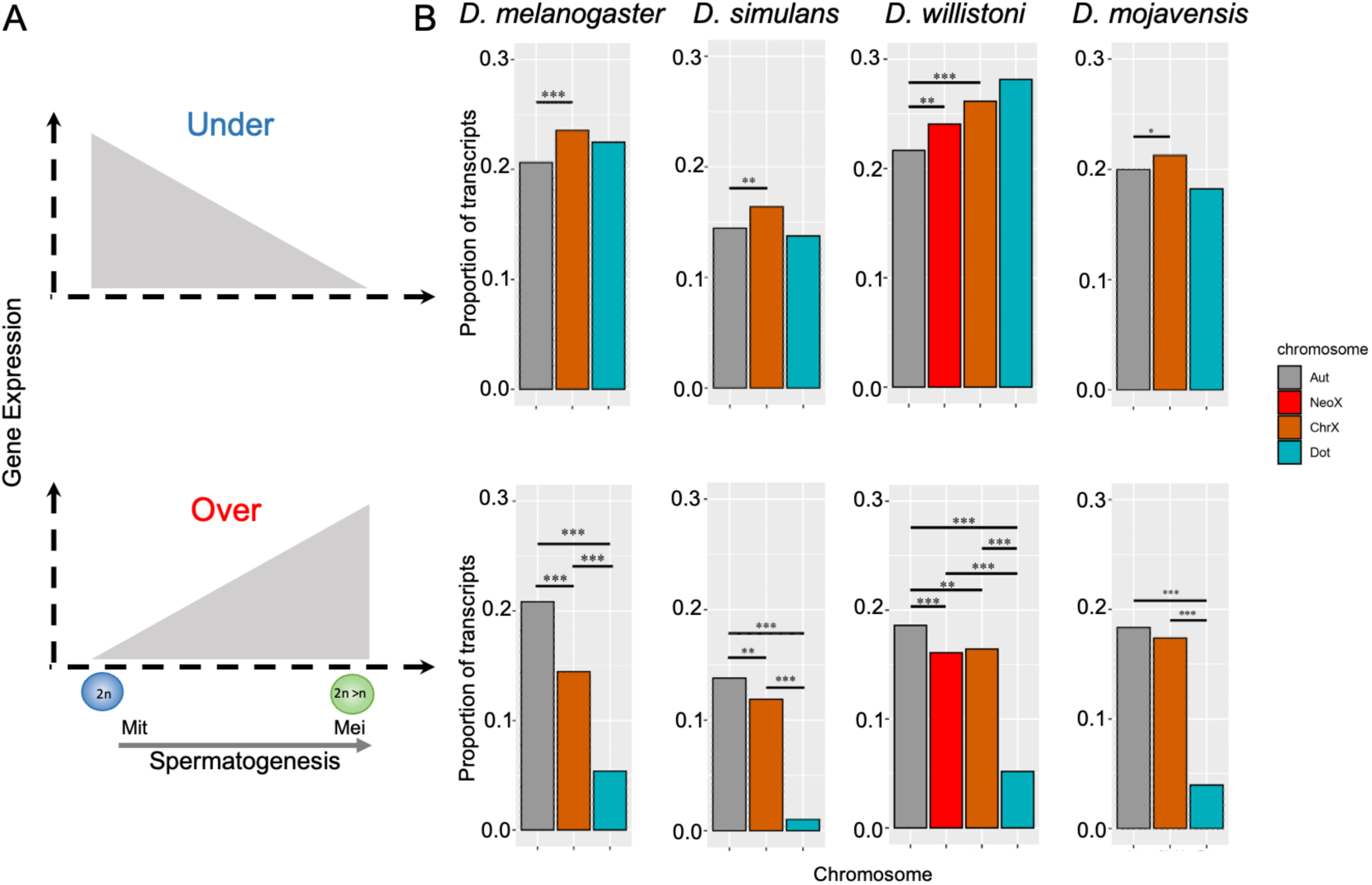
Chromosomal distribution of coding genes over and underexpressed in *Drosophila* male meiosis. **A)** Schematization of meiotic profiles. Under: genes underexpressed in meiosis in relative to mitosis; Over: genes overexpressed in meiosis relative to mitosis **B)** Chromosomal proportion of over and underexpressed transcripts in *D. melanogaster, D. simulans*, *D. willistoni,* and *D. mojavensis*, respectively. Significant proportion differences (Chi-square test with Yates correction) are indicated by *, **, and ***, indicating p values ≤ 0.05, ≤0.01, and ≤0.001, respectively. Autosomes, Neo-X, X and Dot chromosomes are indicated by the color legend by Aut, NeoX, ChrX and Dot, respectively.

Interestingly, the Neo-X chromosome of *D. willistoni* exhibits patterns similar to those of the X chromosome (Figure 2-B). Originating from a fusion and diverging from the *melanogaster* group around 32 million years ago (Tamura et al., 2004), *D. willistoni*’s Neo-X shares key features with the X chromosome, such as demasculinization, indicated by a lower proportion of genes preferentially expressed in males (Sturgill et al., 2007; Vibranovski et al., 2009; Meisel et al., 2009). These results indicate that Neo-X regulation mirrors that of the X chromosome, highlighting the widespread and enduring impact of MSCI. Even though meiosis being the peak of transcriptional activity in *Drosophila* spermatogenesis (White-Cooper, 2009), our findings corroborate an MSCI-consistent reduction in the expression of X-linked and Neo-X genes during meiosis compared to mitosis, implying a robust and ancestral phenomenon, not exclusive to *D. melanogaster* or any specific branch.

A layer of complexity to *Drosophila* sex chromosome evolution is the fourth chromosome, also known as “dot”. Evidence suggests that the dot originated from an ancestral X chromosome and later reverted to an autosome (Vicoso & Bachtrog, 2013; Vicoso & Bachtrog, 2015). However, the dot chromosome does not show the hallmark underexpression linked to inactivation in any species, including *D. melanogaster*, where single-cell studies have noted an overall reduction in expression (Mahadevaraju et al., 2021). Although its significantly lower transcript counts—about 18 times fewer than other chromosomes (Figure 2-B) - might limit the statistical power to detect substantial differences, this low count still reveals enrichment of overexpressed genes on autosomes compared to the dot. This suggests that its MSCI signature, a remnant of the dot chromosome’s X chromosome origin, may be weaker or less conserved across the genus.

### MSCI affects Long non-coding gene expression

To investigate whether the MSCI pattern extends to *Drosophila* lncRNAs — an area not yet explored — we focused our analysis on RNA-seq data from *Drosophila melanogaster* due to its more comprehensive genome annotation (Matthews et al., 2015). The assessment revealed significant enrichment of underexpressed genes during meiosis on the X and dot chromosomes, with autosomes exhibiting a higher proportion of overexpressed genes (Figure 3-B).

**Figure 3.**
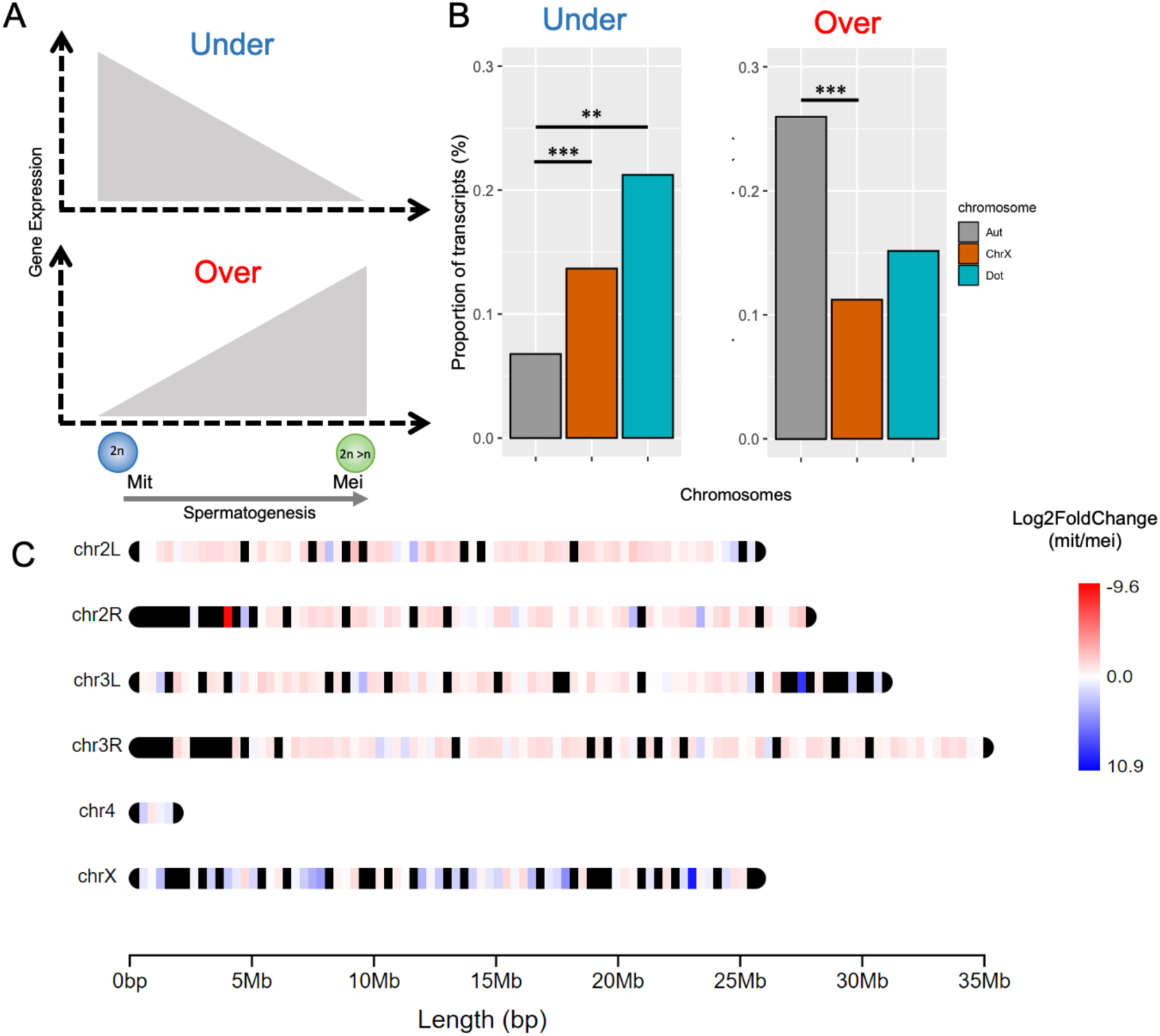
*D. melanogaster* chromosomal proportions of under and overexpressed transcripts of lncRNAs. **A)** Schematization of meiotic profiles. **B)** Chromosomal proportion of over and underexpressed transcripts. **C**) Detailed chromosomal distribution of expression patterns of lncRNAs. Regions in red represent overexpressed genes, while blue represents the opposite pattern. Significant proportion differences (Chi-square test with Yates correction) are indicated by *, **, and ***, indicating p values ≤ 0.05, ≤ 0.01, and ≤ 0.001, respectively. The lncRNAs were analyzed only in *Drosophila melanogaster* due to its extensive genome annotation. Autosomes, X and Dot chromosomes are indicated by the color legend by Aut, ChrX and Dot, respectively.

In mammals, somatic X chromosome inactivation is driven by the long non-coding RNA Xist, which initiates silencing by spreading across the X chromosome from the X inactivation center (XIC) (Lyon, 1961; Brown et al., 1991). Inspired by this spatial regulation mechanism, we sought to understand the distribution of downregulated long noncoding genes on the *Drosophila* X chromosome. To achieve this, we mapped the chromosomal coordinates of each lncRNA, revealing that those differentially expressed during meiosis are uniformly distributed along the X chromosome (Figure 3-C).

Additionally, our RNA-seq results reveal that long noncoding gene expression is predominantly post-meiotic, with 197 transcripts highly expressed during mitosis, 137 during meiosis, and 846 post-meiosis (Table S1). This observation supports previous studies indicating that most lncRNAs likely serve post-meiotic functions (Wen et al., 2016; Vedelek et al., 2018).

### Evolutionary Age-Dependent Gene Expression Dynamics in Spermatogenesis

If MSCI is a widespread evolutionary effect, an intriguing question arises about the fate of genes that have recently originated on the X chromosome. In *D. willistoni*, MSCI appears to be as robust on newly incorporated X-linked regions as it is on the *Drosophila* genus X chromosome (Figure 2-B), suggesting a whole-chromosome response. However, it would also be valuable to explore how MSCI impacts individual genes that arise on the X chromosome. To answer this question, we examined genes that recently emerged on the X chromosome of *D. melanogaster* (Zhang et al. 2010, complete data on new genes are not available for the other species). Figure 4 offers a comprehensive overview of the dynamics of gene expression changes across the stages of spermatogenesis, elucidating the influence of age on these intricate processes.

**Figure 4.**
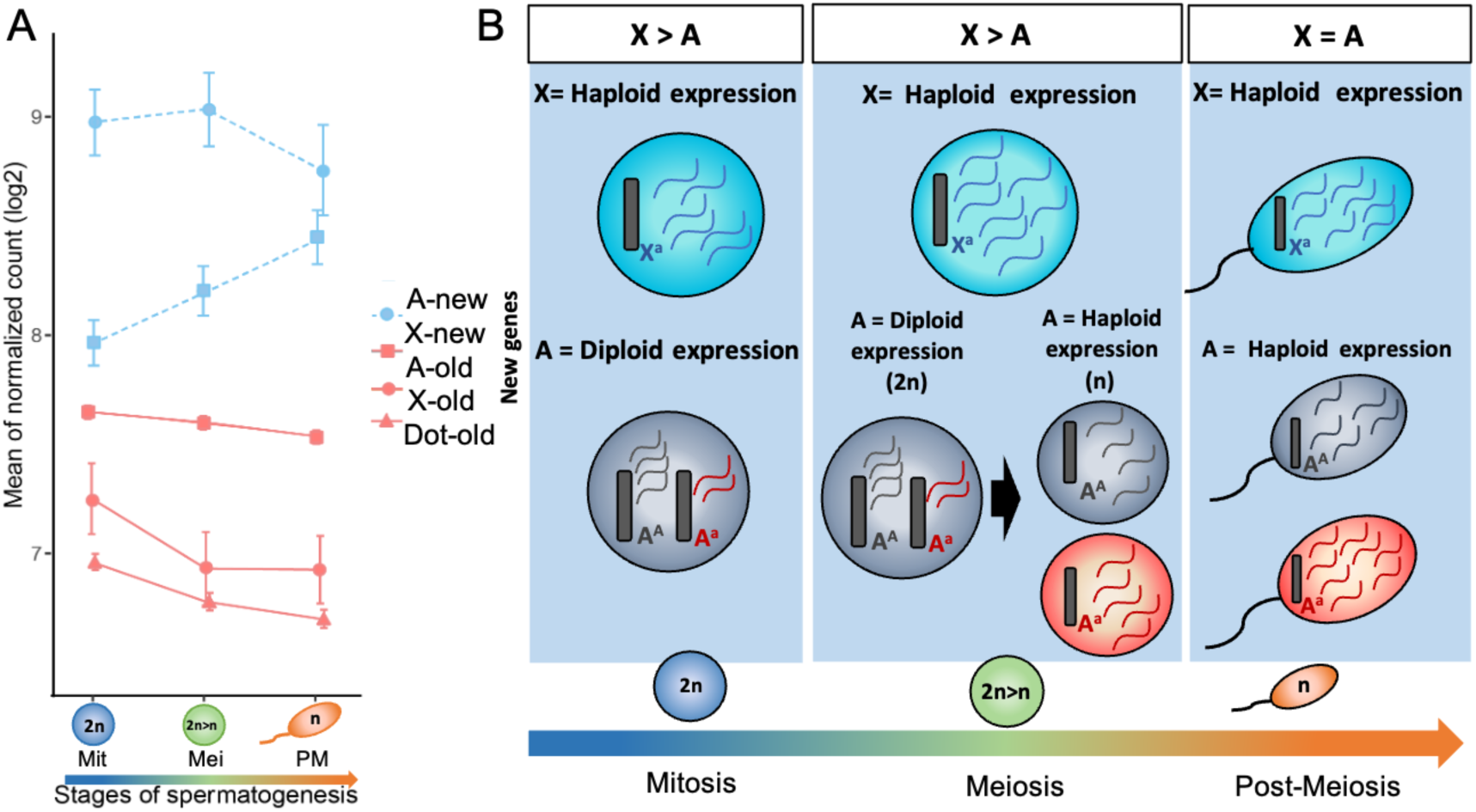
Gene Expression Dynamics by Gene Age and Chromosomal Location. **A)** Average transcript expression (log₂ normalized) across stages of spermatogenesis for newly evolved genes (blue) and older genes (salmon) located on the X chromosome, autosomes, and dot chromosome, denoted by circles, squares, and triangles, respectively. Error bars indicate the standard error of the mean (SEM). Notably, no newly evolved genes were detected on the dot chromosome. **B)** Diagrams illustrating haploid and diploid expression dynamics for new genes on the X chromosome and autosomes. The diagrams depict cell phenotypes based on gene expression relative to chromosomal location and dominance effects: turquoise indicates X-linked allele expression (X^a^), red denotes autosomal recessive allele expression (A^a^), and gray represents autosomal dominant allele expression (A^A^). mRNA expression levels are color-coded accordingly. Note that meiosis involves a diploid phase until the completion of meiosis I, followed by a haploid phase. The top rectangles highlight, for each phase, the relative selective advantage of chromosomal location for an adaptive recessive mutation in a newly evolved gene and, consequently, the expected relative expression.

In the context of *D. melanogaster,* genes that emerged before the split of the *Sophophora* and *Drosophila* subgenera (*i.e.,* old genes) show a progressive decrease in X-linked expression throughout spermatogenesis. This decline, also observed for dot-linked genes, is particularly pronounced from mitosis to meiosis (Figure 4-A, Wilcoxon: p-value < 2.2e-16), coinciding with the stage where X chromosome inactivation occurs. Notably, this phenomenon is absent from the expression of autosomes (Figure 4-A) and persists, at least to some extent, in the post-meiotic stages. Given that old genes constitute over 90% of the total count of *D. melanogaster* transcripts, their expression patterns align with the general MSCI profile observed within this species.

In contrast, newly evolved X genes exhibit a markedly higher expression during both the mitotic and meiotic phases (Wilcoxon: p-value ≤ 0.009), surpassing the mean expression of autosomal genes in these phases (Figure 4-A). Therefore, it is clear that MSCI mechanisms do not influence newly emerged genes, implying that MSCI might take longer to affect individual sequences or may be incapable of extending to new regions once it is established.

Our three-phase spermatogenic profiling enables a detailed examination of gene expression dynamics not only in mitosis and meiosis but also in the post-meiotic phase. Across various organisms—including mammals, plants, and *Drosophila*—new genes, primarily located on autosomes, tend to show heightened expression in the later stages of the male germline (Soumillon et al., 2013; Cui et al., 2015; Raices et al., 2019). This pattern is linked to the advantages of haploid selection in the early evolution of recently emerged genes, as recessive adaptive mutations can be directly exposed to natural selection during this stage (Raices et al., 2019; Figure 4-B). Our RNA-seq analysis supports these findings, showing that, on average, newly evolved autosomal genes display a gradual increase in expression throughout spermatogenesis (Figure 4-A).

A novel insight, however, comes in the post-meiotic phase, where we observe a convergence in mean expression levels between newly evolved autosomal and X-linked genes (Figure 4-A, Wilcoxon: p-value = 0.68). This pattern suggests a broad selective advantage for new genes in haploid expression: while the hemizygous state of X-linked genes offers this benefit consistently across all germline phases, autosomal genes experience this advantage exclusively in the post-meiotic phase (Figure 4-B).

## Discussion

Our data indicates that MSCI was established before the divergence of the *Drosophila* and *Sophophora* subgroups (Figure 5), as genes that are underexpressed during meiosis (relative to mitosis) are significantly more prevalent on the X chromosome than on autosomes across the four analyzed *Drosophila* species. While this finding provides valuable insight into the shared ancestral origin of MSCI within the *Drosophila* genus, it leaves open the question of whether MSCI predates this divergence.

**Figure 5.**
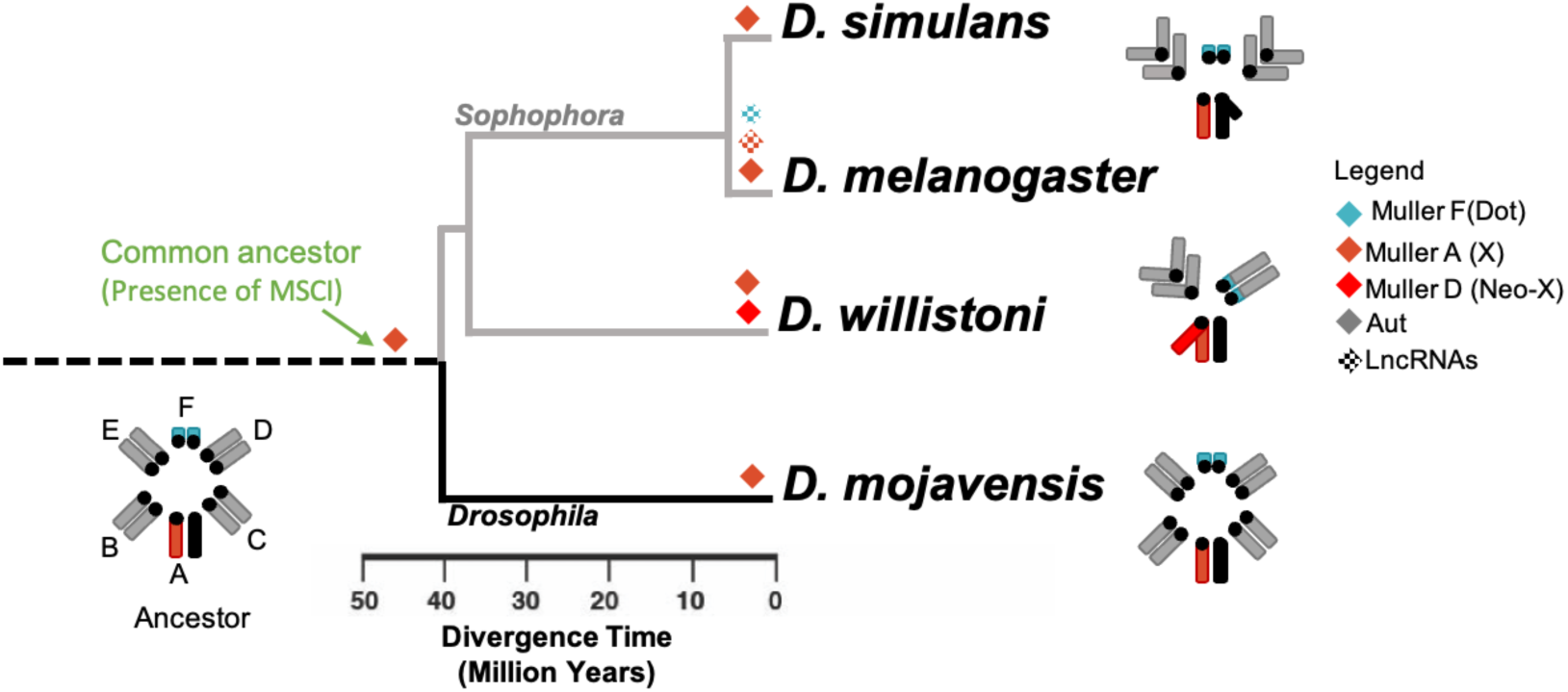
Proposed MSCI evolution in the *Drosophila* genus. The phylogeny is adapted from FlyBase (2023). The green arrow indicates the proposed emergence of Meiotic Sex Chromosome Inactivation (MSCI) within the *Drosophila* lineage. Chromosome morphology transitions are depicted along the tree, with color-coded diamonds indicating the chromosomal elements that experienced downregulation during spermatogenesis.

Indeed, recent single-cell RNA-seq studies have observed X chromosome downregulation during male meiosis in mosquitoes as well (Taxiarchi et al., 2021; Page et al., 2023). Interestingly, the mosquito X chromosome includes genes from the A and F Muller elements found in *Drosophila* (Blackmon et al., 2015; Holt et al., 2002). Further investigation is needed to determine whether MSCI emerged before the evolutionary split between *Drosophila* and mosquitoes or if it developed independently within different Diptera lineages.

Although the dot chromosome (Muller F), as the ancestral X, shares several characteristics with the current X chromosome, it does not appear to be broadly affected by MSCI across the *Drosophila* genus. In *D. melanogaster*, in line with our findings (Figure 4-A), previous studies have shown that dot-linked genes are significantly less expressed in larval testes compared to autosomal genes (Mahadevaraju et al., 2021), a pattern also observed for long non-coding RNAs (Figure 3). However, in other species, the Muller F chromosome consistently shows an underrepresentation of overexpressed genes during meiosis but lacks the significant enrichment of underexpressed genes — a key feature of MSCI (Figure 2-B).

This discrepancy could be due to several methodological, mechanistic, and evolutionary factors. Methodologically, the lower transcript counts on the dot chromosome may reduce the power to detect a full MSCI effect, particularly in non-*melanogaster* species with less refined genome annotations. Mechanistically, studies in *D. melanogaster* have shown that the dot chromosome is frequently near, but not entirely adjacent to, the RNA polymerase-inactive X chromosome territory in spermatocytes, possibly leading to a partial or weaker MSCI effect (Mahadevaraju et al., 2021). Evolutionarily, the ancestral X chromosome has been diploid throughout *Drosophila*’s history, which may have lessened the selective pressures associated with a lack of pairing, potentially weakening MSCI constraints over time (Riddle & Elgin, 2018).

The advantage of our cell type enrichment dissection method over the testis single-cell RNA-seq approach lies in its capacity to profile the expression of post-meiotic cells. Single-cell RNA-seq encounters difficulties in sequencing non-round cells, leading to a reduction in the number of cells available from the post-meiotic stage, particularly elongated and mature spermatids corresponding to 5.82% of the total number of cells from spermatocyte cluster (1435 late spermatocytes and 83 late spermatids, from Witt et al., 2019). This limitation arises from the necessity to remove the sperm tail as part of the technique, a process that introduces stress to the cells and has the potential to alter their transcriptional profile (Li et al., 2021).

Our latest RNA-seq findings throughout spermatogenesis provide new insights, showing that newly formed X-linked genes consistently exhibit higher expression across all phases. This discrepancy with previous microarray data may be due to RNA-seq’s increased sensitivity and the improved annotation of the *Drosophila melanogaster* genome, which together allow for the detection of a broader range of transcripts. The elevated expression, particularly during meiosis, confirms that new X-linked genes may take time to become fully subject to MSCI effects (Raices et al. 2019).

This sustained expression across mitosis, meiosis, and post-meiosis aligns with the concept of haploid selection and supports the Faster-X hypothesis, which suggests that the hemizygous state of the X chromosome in males accelerates the evolution of X-linked genes compared to autosomes (Charlesworth et al., 1987). In contrast, autosomal genes only expose recessive adaptive mutations to selection during the haploid stages (Immler, 2019). As a result, the expression boost driven by haploid selection is primarily seen at the end of meiosis and into post-meiosis, a phenomenon unique to autosomal genes. This allows newly evolved autosomal genes, driven by positive selection, to reach expression levels comparable to new X-linked genes, though without surpassing them.

Importantly, our study has provided these insights, enhancing not only our understanding of MSCI evolution within the *Drosophila* genus but also enriching the broader discourse on the evolution of newly formed genes on the X chromosome. These findings contribute to a deeper comprehension of the genetic and evolutionary mechanisms underlying spermatogenesis and gene regulation, offering new perspectives on the dynamics of genome evolution.

## Materials and Methods

### Fly stocks and sample preparation

By incorporating species with diverse evolutionary relationships and karyotypes, along with maintaining comparable testis morphology, the study aims to uncover evolutionary patterns in sex chromosome expression regulation while controlling for genetic and morphological variations. The *Drosophila* testis, exhibiting variations in size across the studied species, consistently maintains a characteristic shape among the selected ones: a long, coiled tube, presenting a thin and elongated structure (Schärer et al., 2008). The similarity in the tubular shape among these species reflects commonalities in how spermatogenic developmental cells are arranged sequentially, extending from the apical to the distal end (Figure 1-A). *Drosophila* spermatogenesis starts in the apical end, which is enriched with stem hub cells, spermatogonial stem cells, and interconnected spermatogonia, corresponding to the mitotic cells (Fuller, 1993). The spermatogonial cells undergo four rounds of mitotic divisions, and after completing the premeiotic S-phase, they undergo enlargement and extensive transcriptional changes, generating cysts with 16 primary spermatocytes. At this point, they are concentrated in the proximal region and their volume increases by 20-25 times, followed by cysts with 32 secondary spermatocytes. The distal region is enriched with post-meiotic cells, such as 64 round spermatid cysts, elongated spermatids, and spermatozoa (Cenci et al., 1994; Fuller, 1993). Samples of the different phases of spermatogenesis (mitosis, meiosis, and post-meiosis) were prepared following the methods outlined in Vibranovski et al. (2009) for *D. melanogaster*, with minor adjustments for the other species regarding their cell distribution (see Figure S1). Briefly, we dissected testes (without seminal vesicles) in PBS from virgin males aged 6-10 days after eclosion, using 0.25 mm diameter insect pins. Fifty to one hundred dissections per sample were isolated and stored in RNAlater® at - 20 °C until extraction.

### RNA isolation, RNA-seq library generation, and Quality Control

Total RNA was extracted using PicoPure™ RNA Isolation Kit (Arcturus). Until the sequencing and quality control, RNA samples were stored in RNAstable® (Biomatrica). RNA hydration, quality control tests, library preparation, and sequencing were done at the Genomics Facility at the University of Chicago. Quality control of the samples was carried out on a Bioanalyzer 2100 instrument (Agilent). Stranded multiplexed libraries were sequenced on the HiSeq Illumina 4000 platform. The *D. melanogaster* libraries were sequenced using a 50 bp single-end protocol while the remaining species were sequenced using 100 bp paired-end. Each combination of species and spermatogenesis phase was done with at least three biological replicates and 30 million fragments per replicate.

### Expression analyses and Data Quality Control

In all four cases, we included coding (CDS) and non-coding sequences (lncRNA). For *D. melanogaster*, we used the reference genome Release 6.34 (ref FlyBase, May 2020). For the other three species, (*D. willistoni, D. mojavensis*, and *D. simulans*), we started from NCBI RefSeq releases 101, 101, and 103 respectively (NCBI currently serves as the primary platform for genome and annotation updates for these species). Notably, for the Y chromosome, the original sequences from the designated releases were substituted with sequences identified by Vanderlinde, 2016 for *D. mojavensis*, and Ricchio et al., 2021 for *D. willistoni*. To refine the dataset, CD-HIT (version: 4.8.1) was employed to eliminate redundant sequences, following the methodology by Li and Godzik (2006) and Fu et al. (2012).

FastQ reads were aligned to the corresponding reference transcriptome with the RSEM pipeline (Li & Dewey, 2011) using bowtie2 (Langmead & Salzberg, 2012) as the aligner. Estimated counts were imported to R with Tximport (Soneson, Love & Robinson, 2015) for differential expression analysis with DeSeq2 (Love; Anders & Huber, 2014), which employs paired Wald tests with false discovery rate (FDR) correction (Benjamini & Hochberg, 1995). Pearson correlations were estimated using log2 transformations (Figure S3). To identify different spermatogenic expression profiles, K-means clustering was applied to all transcriptomes, including the previous microarray-generated database for *D. melanogaster* (Vibranovski et al. 2009).

Chromosomes’ locations were collected from the reference annotations for *D. melanogaster* and *D. simulans*. For *D. willistoni* and *D.mojavensis*, scaffolds were mapped to chromosomes using genome coordinates according to Schaeffer et al., 2008 (tables S5, S6). The ancestral dot chromosome was fused to the Muller E element in the *D. willistoni* lineage (Papaceit & Juan, 1998; Schaeffer et al., 2008). For this species, we used the mapping information provided by Schaeffer et al., 2008, in which genes proximal to CG17119-PA (E), were considered as dot genes (scaffold scf2_1100000004943. See Figure S6). Ortholog relationships among the genes of the four species were retrieved from FlyBase (2020).

### List of new genes identification

The dataset of new genes in *Drosophila* was obtained from the GenTree platform (http://gentree.ioz.ac.cn/), (Shao et al., 2019). These genes are categorized into six age groups based on their age (Zhang et al., 2010). The youngest group is exclusive to *Drosophila melanogaster* and is labeled as branch 6, with subsequent branches denoted accordingly. We grouped branches 1-6, corresponding to new genes (less than 62 million years old), and Branch 0, which comprises orthologs found both in the S*ophophora* and *Drosophila* subgenus, forming the old genes group (Table S1).

## Data availability

RNA-Seq reads were uploaded to NCBI under the BioProject accession ID: PRJNA1188117.

## Supporting information

Supporting informations

## Acknowledgments

We thank members of our labs and colleagues for insightful discussions. We also thank Denise Selivon Scheepmaker for the primers used for *Wolbachia* detection.

## Author Contribution

Conceptualization: C.C.A; M.D.V. Methodology: C.C.A; C.A.M; M.D.V; A.B.C. Investigation: C.C.A; C.A.M; M.D.V; G.G. Supervisions: M.D.V; A.B.C. Writing-original draft: C.C.A; M.D.V. Writing-review and editing: C.C.A; M.D.V; C.A.M; A.B.C; G.G

