## Supporting informations for "Meiotic Sex Chromosome Inactivation: conservation across the *Drosophila* genus"

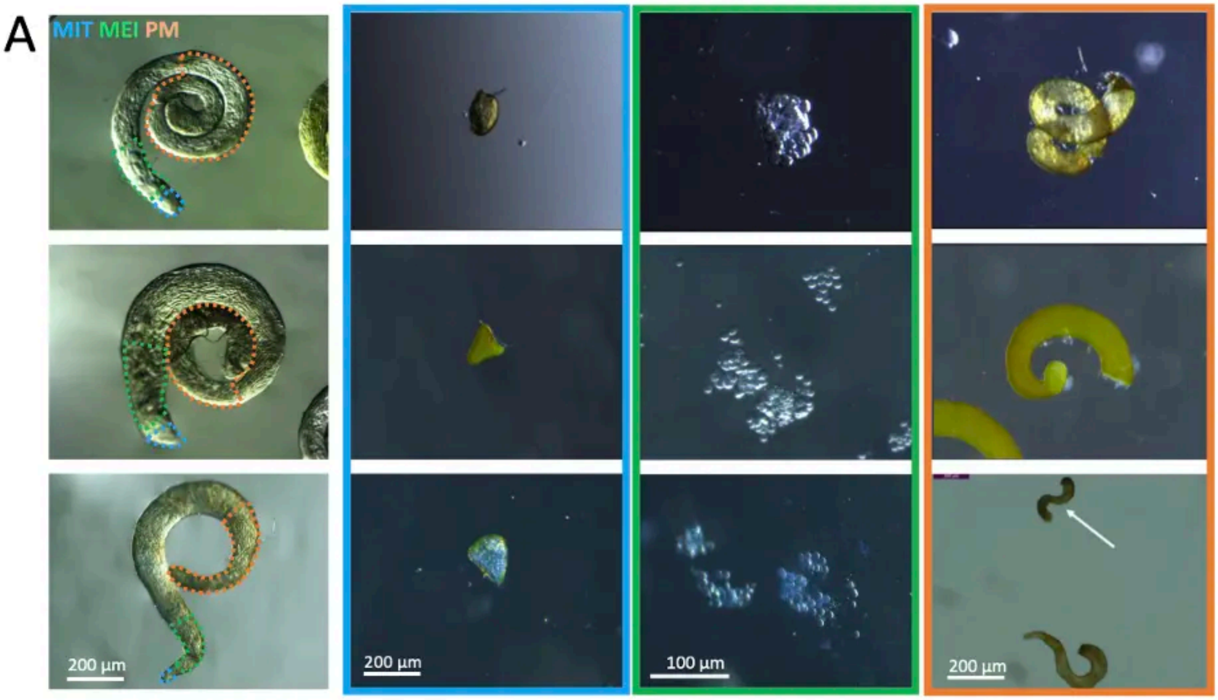


B

| Specie | Mitosis | Meiosis | Post Meiosis |
|---|---|---|---|
| *D. melanogaster* | 110 | 50 | 100 |
| *D. simulans* | 50 | 54 | 46 |
| *D.willistoni* | 80 | 57 | 38 |
| *D.mojavensis* | 90 | 68 | 45 |

**Figure S1. Stage-Specific Testis Dissection. A)** Testis regions corresponding to the three primary phases of spermatogenesis in *D. melanogaster/D. simulans* (first row), *D. mojavensis* (second row), and *D. willistoni* (third row). Each species displays the entire testis, with specific regions enriched with specific cell types highlighted: blue areas indicate mitotic cells, green areas indicate meiotic cells and orange-dotted areas indicate post-meiotic cells. The proportions of these regions vary among species. orange-dotted areas indicate post-meiotic cells, green areas indicate meiotic cells and blue areas indicate mitotic cells. The proportions of these regions differ between species, with arrows marking the distal region containing post-meiotic cells. **B)** Average sample size for each dissected testis region across the species.

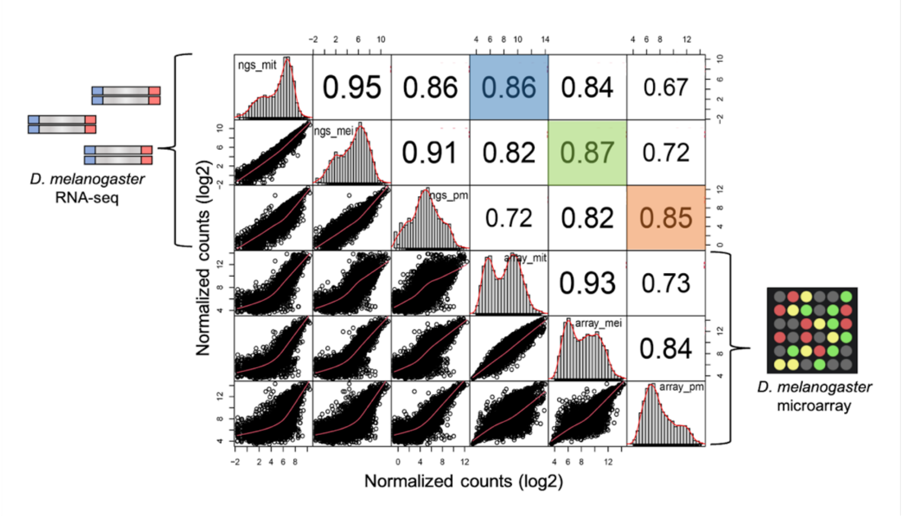


**Figure S2. Pearson Correlation of Gene Expression in *Drosophila melanogaster* across Different Developmental Stages and Techniques.** Pairwise comparisons of gene expression data in *Drosophila melanogaster* across three developmental stages (mitotic, meiotic, and post-meiotic) using two different techniques: RNA sequencing (NGS) and microarray analysis (Array) from Vibranovski *et al*. (2009). The colors represent the correlation coefficients for stage-specific data, with blue, green, and orange corresponding to Mitosis (mit), Meiosis (mei), and Post-Meiosis (pm), respectively. The upper panel displays the pairwise correlation coefficients, while the lower panel shows scatter plots with normalized counts. The diagonal histogram illustrates the distribution of gene expression for each developmental stage in each technique.

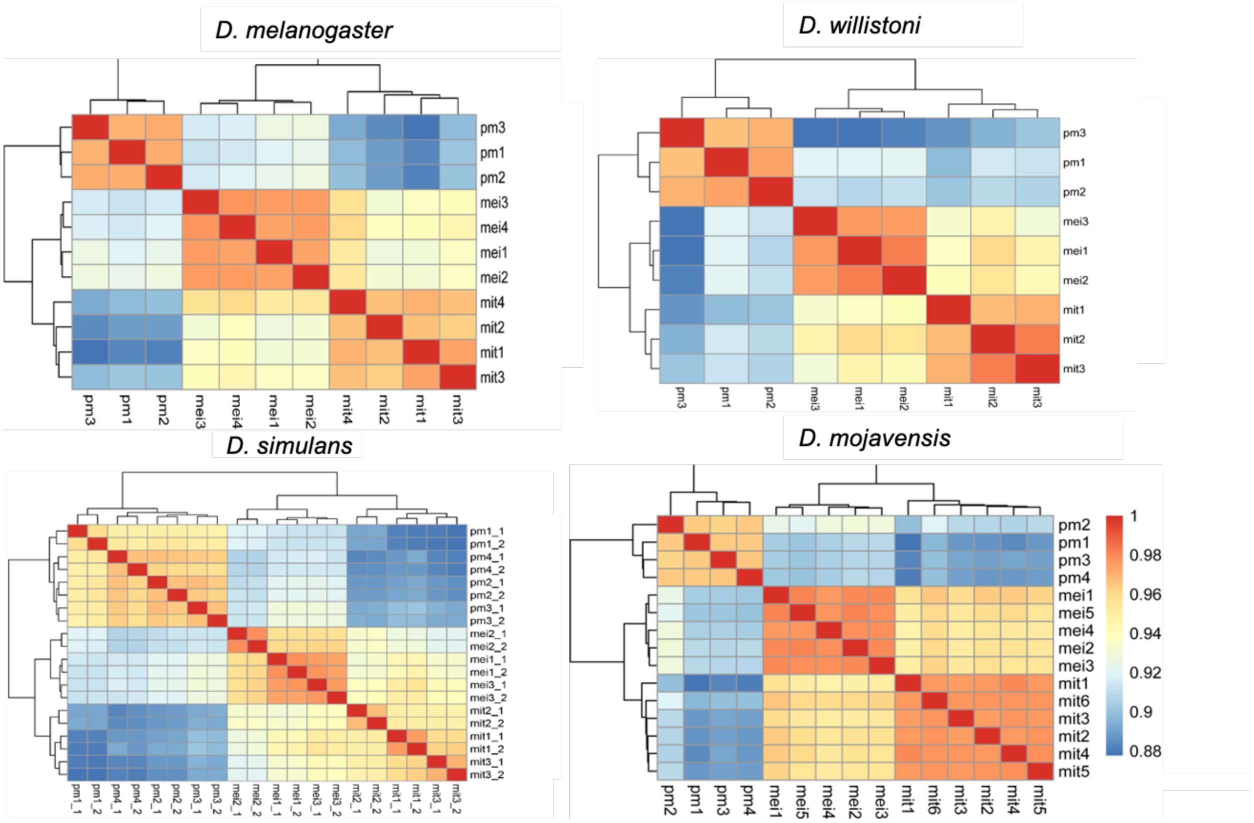


**Figure S3. Similarity analyses among biological and technical replicates spermatogenic phases for *Drosophila* species**. Heatmaps evaluate similarities among mitosis (mit), meiosis (mei), and post-meiosis (pm)

and within biological replicates, with warmer tones indicating higher Pearson correlations. In *D. simulans*, _1 and _2 represent the technical replicates.

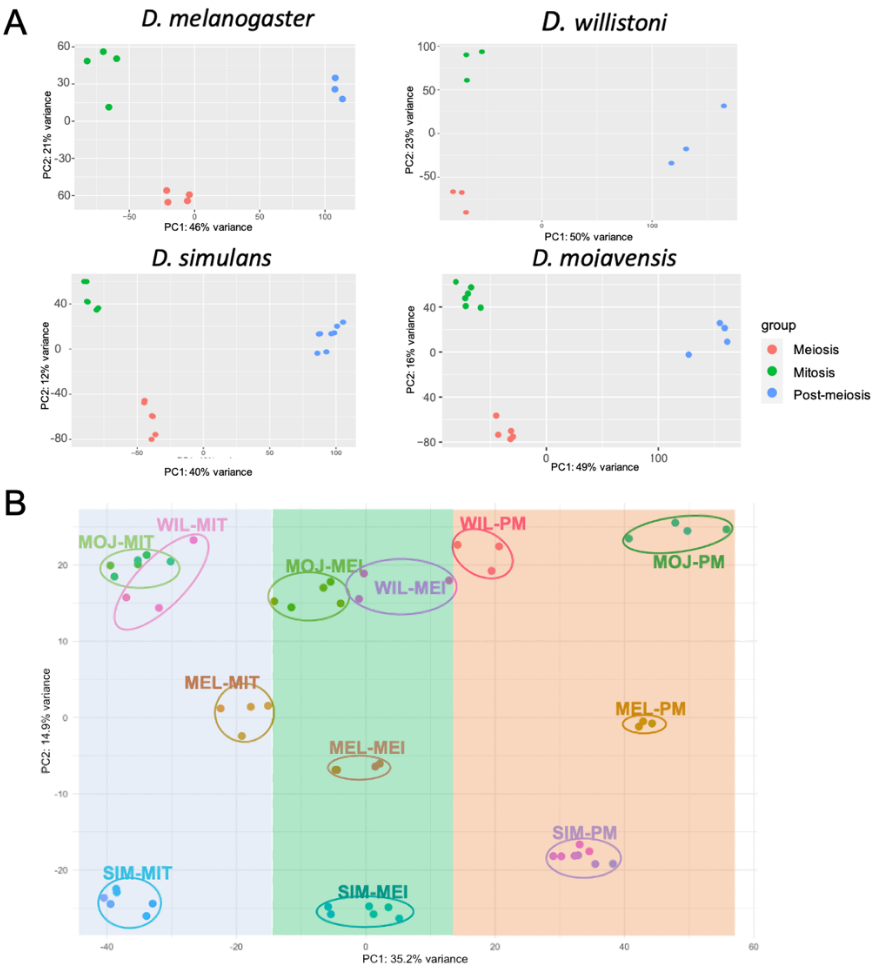


**Figure S4. Principal component analysis (PCA) of intra and inter-species sample replicates. A)** PCA plots showing the relationships among spermatogenesis stages (mitosis, meiosis, post-meiosis) within each species. Stages are color-coded: mitosis (green), meiosis (orange), and post-meiosis (blue), highlighting distinct expression profiles associated with each stage in *D. melanogaster*, *D. simulans*, *D. mojavensis*, and *D. willistoni*. **B)** PCA of orthologous genes shared across all species, grouping samples by spermatogenesis stages across species. Colored rectangles on the background indicate mitotic (blue), meiotic (green), and post-meiotic (orange) samples. The first principal component (PC1) explains 35.2% of the variance, separating stages of spermatogenesis, while the second component (PC2) accounts for 14.9% of the variance, distinguishing species within the same stage. This pattern highlights a higher similarity in gene expression within stages across species than between stages, underscoring the conserved nature of spermatogenesis gene expression across the *Drosophila* genus.

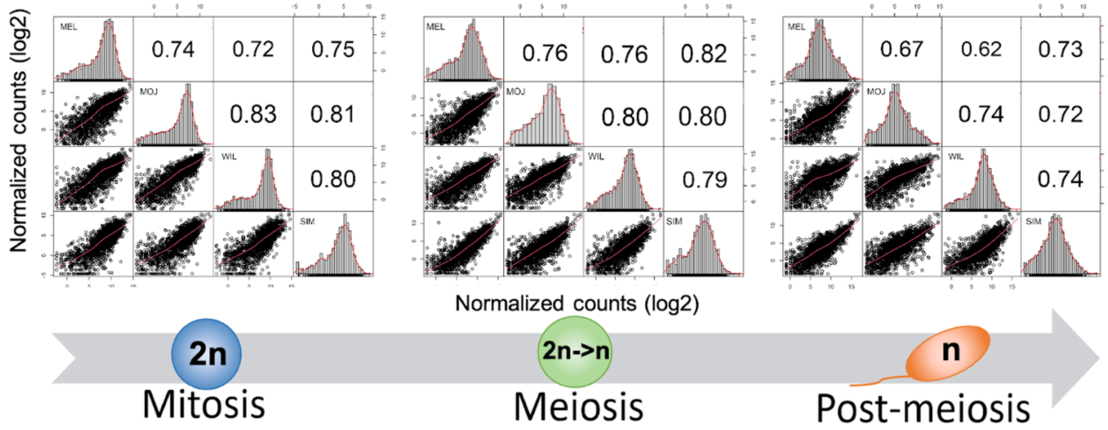


**Figure S5. Pairwise Correlations of Gene Expression.** This figure presents pairwise Pearson correlations between orthologous genes in *Drosophila* species during spermatogenesis profiles for *D. melanogaster* (MEL), *D. mojavensis* (MOJ), *D. willistoni* (WIL), and *D. simulans* (SIM), with a sample size of 1913 genes. The upper panel displays the pairwise correlation coefficients, while the lower panel shows scatter plots with normalized counts. The diagonal histogram illustrates the distribution of gene expression for each species at each stage.

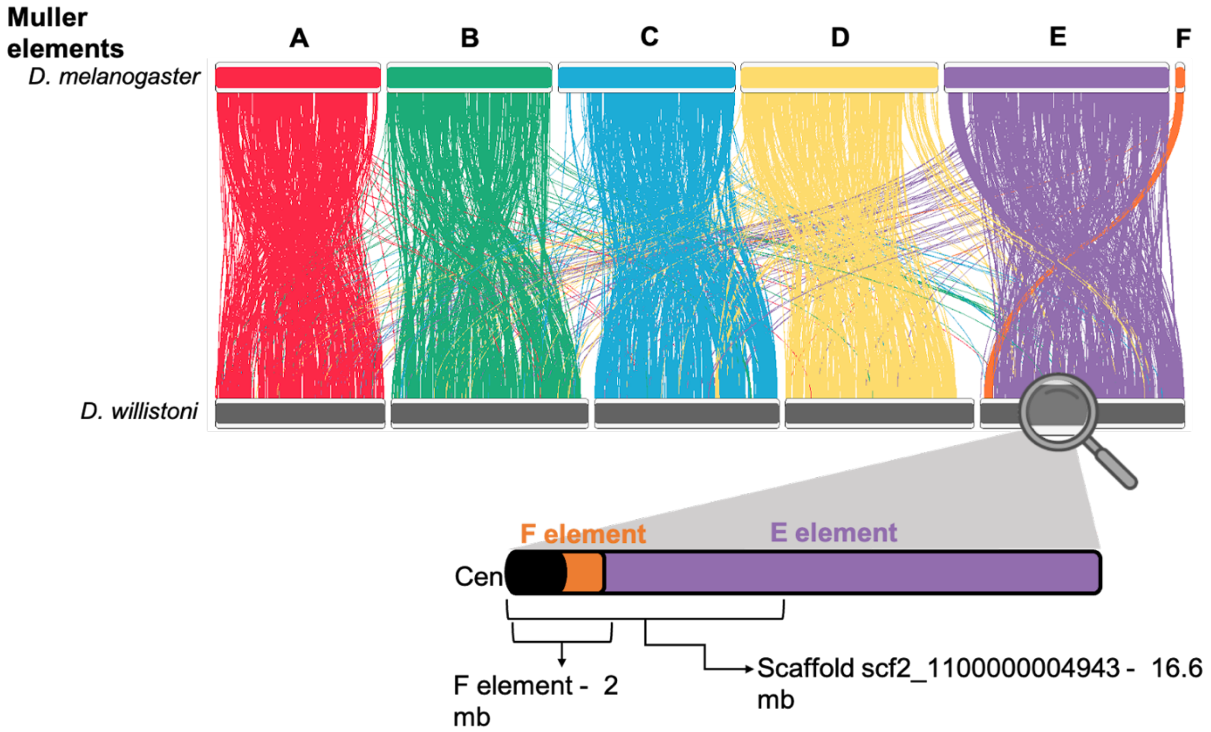


**Figure S6. Muller E-F Mapping in *D. willistoni*.** The graph illustrates chromosome synteny between *D. melanogaster* and *D. willistoni*, based on Schaeffer et al. (2008). Each bar represents a Muller Element, visualizing the alignment using RIdeogram (PMID: 33816903). The lower panel zooms in on the E element in *D. willistoni*, highlighting the portion corresponding to the F element, also known as the dot chromosome. The dot chromosome corresponds to chromosome section 78A-78D. The junction between the E and F elements is defined by the genes CG34036-PA (F) and CG17119-PA (E), and the region spans nucleotides 2,014,728 to 2,029,101, as described by Papaceit & Juan (1998) and Schaeffer et al. (2008). "Cen" indicates the centromere.
